# DNA methylome combined with chromosome cluster-oriented analysis provides an early signature for cutaneous melanoma aggressiveness

**DOI:** 10.1101/2022.04.11.487909

**Authors:** Arnaud Carrier, Cécile Desjobert, Loïc Ponger, Laurence Lamant, Matias Bustos, Jorge Torres-Ferreira, Rui Henrique, Carmen Jeronimo, Luisa Lanfrancone, Audrey Delmas, Gilles Favre, Antoine Daunay, Florence Busato, Dave S.B. Hoon, Jörg Tost, Chantal Etievant, Joëlle Riond, Paola B. Arimondo

## Abstract

Aberrant DNA methylation is a well-known feature of tumours and has been associated with metastatic melanoma. However, since melanoma cells are highly heterogeneous, it has been challenging to use affected genes to predict tumour aggressiveness, metastatic evolution, and patients’ outcomes. We hypothesized that common aggressive hypermethylation signatures should emerge early in tumorigenesis and should be shared in aggressive cells, independent of the physiological context under which this trait arises.

We compared paired melanoma cell lines with the following properties: *(i)* each pair comprises one aggressive counterpart and its parental cell line, and *(ii)* the aggressive cell lines were each obtained from different host and their environment (human, rat, and mouse), though starting from the same parent cell line. Next, we developed a multi-step genomic pipeline that combines the DNA methylome profile with a chromosome cluster-oriented analysis.

A total of 229 differentially hypermethylated genes were commonly found in the aggressive cell lines. Genome localization analysis revealed hypermethylation peaks and clusters, identifying eight hypermethylated gene promoters for validation in tissues from melanoma patients.

Five CpG identified in primary melanoma tissues were transformed into a DNA methylation score that can predict survival (Log-rank test, *p*=0.0008). This strategy is potentially universally applicable to other diseases involving DNA methylation alterations.

## INTRODUCTION

Cutaneous metastatic melanoma is the deadliest form of skin cancer and its occurrence is growing (1). The recent development of targeted and immune therapies has dramatically improved patient’s outcomes. Indeed, median overall survival of patients with advanced-stage melanoma has increased from ∼ 9 months to at least 2 years since 2011 (2). Overall survival is better after targeted (3) or immuno therapies (4), but there are still non-responders and neo/acquired resistants. Despite these advances, there is place for improvement in particular to discover novel early prognostic markers and potential avenue for adjuvant therapies.

DNA methylation in malignant melanoma has been studied to identify specific DNA methylation changes and decipher their impact. Melanoma has a CpG island methylator phenotype (CIMP) (5), and several methylated genes are associated with melanoma progression (6), with aggressive clinical and pathological features and poor survival in patients (7, 8), are candidate epigenetic drivers of melanoma metastasis (9) or are implicated in immunotherapy resistance (10). Such DNA methylation changes have been studied at different stages of the metastatic disease, but not in primary cutaneous tumour. Importantly, DNA methylation has been shown to occur very early in tumour formation (11), and thus has the potential to provide early biomarkers indicating the metastatic potential of the tumour. However, the field currently lacks a genomic strategy that can both account for genetic environment and identify early DNA methylation markers that predict the aggressiveness of melanoma.

Here, we developed a strategy that leverages the DNA methylome from different pairs of human melanoma cells lines. Cells within pairs share a common genetic background, but one counterpart has been selected for aggressiveness in different *in vivo* contexts (human *vs* murine). We proposed that the DNA methylation signature of tumour aggressiveness would be independent of the physiological context: starting from a human tumour, shared signatures relevant to aggressiveness should emerge independent on whether this trait were acquired in humans, or whether cells have been implanted into rats or mice. In a multi-step selection process, we identified hypermethylated sites common to the most aggressive melanoma forms, analysed the distribution of these sites in the genome, and validated these methylation peaks in cell lines and patient samples. This strategy identified a DNA methylation signature of five CpG sites in four gene promoters in primary tumours that could predict the overall survival of the patients and thus has potential diagnostic application. This strategy, which overcomes heterogeneity in tumours due to the environment, can potentially be generalized to other cancers involving DNA methylation alterations.

## MATERIAL AND METHODS

### Cell Culture

The WM115 and WM266-4 cell lines were obtained from the American Type Culture Collection. The WM-115 cell line was derived from a human primary melanoma in early stages of the vertical growth phase (VGP). The WM266-4 cell line was derived from a cutaneous metastatic melanoma tumour of the same patient. The WM983A and WM983B cell lines were obtained from the Coriell Institute (USA). The WM983A was derived from a human primary melanoma in VGP and the WM983B is the cutaneous lymph node metastasis from the same patient. The M4Be cell line was established from a human cutaneous lymph node metastasis (12). The TW12 cell line is an aggressive variant of the parental M4Be cell line that was obtained *in vivo* after two serial transplantations (subcutaneous xenografts in new-born immuno-deprived rats). A subclone (TW12) was selected after limiting dilution for its high ability to form lung metastasis (13–15). The M4BeS2 cell line was obtained in the L. Lamant’s laboratory according to the i*n vivo* selection scheme described by Clark et al. (16). Briefly, M4Be cells were xenografted intravenously in nude mice. Lung metastases were collected, grown briefly *in vitro* and used for a second cycle of intravenous injection. Lung metastases were collected and established *in vitro* as the M4BeS2 cell line. WM983A and WM983B cell lines were grown in 20% Leibovitz L-15 medium (v/v), 2% FBS heat inactivated (v/v), 5 μg/mL insulin and 1.68 mM CaCl_2_. All other cell lines were grown in DMEM (Invitrogen, France) supplemented with 10% foetal bovine serum (Sigma, France), 2 mM glutamine, 100 UI/mL penicillin-streptomycin and 1.25 µg/mL fungizone (Invitrogen) in 5% CO_2_. Quantitation of viable cells was performed using an Automated Cell Viability Analyzer (Beckman Coulter Vi-Cell). All cell lines were stored in ampoules in liquid nitrogen after receipt. All cell lines were regularly verified for mycoplasma contamination using the MycoAlert^TM^ Mycoplasma Detection Kit (Lonza, Switzerland) and kept for a limited number of passages in culture.

### Tumour samples

Tumour samples from melanoma patients were obtained from the tumour tissue bank at the Department of Pathology, IUCT-O Toulouse Hospital (France). The study was carried out in accordance with the institutional review board-approved protocols (CRB, AC-2013-1955) and the procedures followed were in accordance with the Helsinki Declaration. Pathological specimens consisted of primary melanomas (*n* = 12), lymph node metastases (*n* = 7), and cutaneous metastases (*n* = 3). Additional primary melanoma (*n* = 5) frozen samples were provided by the Department of Experimental Oncology, European Institute of Oncology, Milan (Italy). The Saint John’ Cancer Institute (formerly John Wayne Cancer institute (USA)) Formalin-fixed paraffin-embedded (FFPE) specimen cohort included tissues including primary melanomas (*n* = 12). A total of 20 FFPE tissues samples from patients diagnosed with cutaneous melanoma between 2007 and 2017 at the Portuguese Oncology Institute of Porto (IPO-Porto) without any neoadjuvant treatment were included in this study. All samples were archived at the Department of Pathology of IPO-Porto. All cases were reviewed by an experienced pathologist and staged according to the 8^th^ edition American Joint Committee on Cancer (AJCC) system (17). Relevant clinical data was collected from medical charts. For DNA extraction, a 4 μm section was cut from a representative tissue block and stained with hematoxylin-eosin. Tumour areas containing >70% transformed cells were delimited, enabling macrodissection in eight consecutive 8μm sections. This study was approved by the institutional ethics committee of IPO Porto (CES-IPOP-FG13/2016). Anonymized clinical information for all the melanoma patients analysed is available, and the clinical pathological features of primary melanoma patients are indicated below (Table1).

**Table 1.** Clinical pathological features of primary melanoma patients

| **Variables** | **n (%)** |
| --- | --- |
| **Mean age (sd)** | 66.40 (15.88) |
| **Gender** |  |
| Male | 26 (53) |
| Female | 23 (47) |
| **AJCC 8<sup>th</sup> stages** |  |
| IV | 12 (24) |
| III | 12 (24) |
| IIIA | 1 (2) |
| IIIB | 2 (4) |
| IIIC | 2 (4) |
| II | 2 (4) |
| IIA | 3 (6) |
| IIB | 6 (12) |
| IIC | 6 (12) |
| IB | 2 (4) |
| unknown | 1 (2) |
| **Mutations** |  |
| *NRAS* | 1 (2) |
| *BRAF* | 6 (12) |
| unknown | 42 (86) |

### Genomic DNA isolation

Genomic DNA from cell lines was performed using the DNeasy Tissue kit (Qiagen, France).

Genomic DNA from frozen patient samples was isolated using the QiaAmp kit (Qiagen, France). DNA extraction from FFPE sections was performed using the FFPE RNA/DNA Purification Plus Kit (Norgen Biotek, Thorold, Canada) in accordance with manufacturer’s instructions. DNA concentration and purity were determined using the NanoDrop Lite spectrophotometer (NanoDrop Technologies, Wilmington, DE, USA).

### Illumina methylation 450K microarray analysis

Genome-wide DNA methylation analysis was performed on three independent samples from each cell line. One microgram of DNA was bisulfite-treated using the EpiTect 96 Bisulfite Kit (Qiagen GmbH, Germany). 200 ng of bisulfite-treated DNA was analysed using Infinium HumanMethylation 450K BeadChips (Illumina Inc., CA, USA). The array allows the interrogation of more than 485,000 methylation CpG sites per sample covering 99% of RefSeq genes, with an average of 17 CpG sites per gene region distributed across the promoter, 5’-UTR, first exon, gene body, and 3’-UTR.

The samples were processed according to the manufacturer’s protocol at the genotyping facility of the Centre National de Génotypage (Evry, France) without any modification to the protocol. We used the GenomeStudio® software (version 2011.1; Illumina Inc.) for the extraction of DNA methylation signals from scanned arrays (methylation module version 1.9.0, Illumina Inc.). Methylation data were extracted as raw signals with no background subtraction or data normalization. The obtained ‘ß’ values – that is, the methylation scores for each CpG range from 0 (unmethylated, U) to 1 (fully methylated, M) on a continuous scale – were calculated from the intensity of the M and U alleles as the ratio of fluorescent signals (ß = (Max(M,0))/ (Max(M,0) + Max(U,0) + 100).

All pre-processing, correction and normalization steps were performed using an improved version of the in-house developed pipeline using subset quantile normalization based on the relation to sequence annotation provided by Illumina (18). Probes were considered as differentially methylated if the absolute value of the difference between robust median ß- values in samples of each phenotypes was higher than 0.2: median cell line 1(ß_1_, ß_2_, ß_3_) – median cell line 2 (ß_1_, ß_2_, ß_3_) ≥ 0.2, where ß_1_, ß_2_, and ß_3_ corresponds to the ß-values in three replicates within each cell line, all with a detection p-value < 0.01. This 0.2 threshold, representing approximately a difference in DNA methylation levels of 20%, corresponds to the recommended differences between samples analysed with the Illumina methylation Infinium technology that can be detected with a 99% confidence.

Differential DNA methylation markers were identified using a combination of two approaches. The performance of individual CpGs was assessed testing the absolute DNA methylation difference between samples of the two phenotypes of interest with different thresholds and permitting a small number of misclassifications. At the same time a vector quantization method (nearest centroid classifier) was used to define CpGs that separate, at a given threshold, the two phenotypes of interest. CpGs that were significant in both tests, were used to calculate a vector using a directed z-score, which was subsequently used to assign new samples to their phenotypic group.

The corresponding genes were obtained from a list of differentially methylated probes using the Illumina annotation file and overlap between gene lists from the three cellular pairs was determined.

The promoter methylation scores (%) reported in Figure S1 were defined as follows: mean (probe1 ß-value to probe n ß-value) x100, where probe 1 to probe n are probes that are differentially methylated between WM266 *vs* WM115 cells (with a difference threshold 0.2 on a scale from 0 to 1) and located in the promoter region (TSS1500-TSS200-5’UTR-1^st^ exon).

### Bisulfite pyrosequencing

Quantitative DNA methylation analysis was performed by pyrosequencing of bisulfite-treated DNA as described in Tost and Gut, 2007 (19). CpGs for validation were amplified using 20 ng of bisulfite treated human genomic DNA and 5–7.5 pmol of forward and reverse primer, one of them being biotinylated. Oligonucleotide sequences for PCR amplification and pyrosequencing are given in the supplementary data (Supplementary Dataset2). Reaction conditions were 1 × HotStar® Taq buffer (Qiagen) supplemented with 1.6 mM MgCl_2_, 100 μM dNTPs and 2.0 U HotStar Taq polymerase (Qiagen) in a 25 μL volume. The PCR program consisted of a denaturing step of 15 min at 95°C, followed by 50 cycles of 30 s at 95°C, 30 s at the respective annealing temperature and 20 s at 72°C, with a final extension of 5 min at 72°C. A total of 10 μL of PCR product was rendered single-stranded as previously described and 4 pmol of the respective sequencing primers were used for analysis. Quantitative DNA methylation analysis was carried out on a PSQ 96MD system with the PyroGold SQA Reagent Kit (Qiagen) and results were analysed using the PyroMark software (V.1.0, Qiagen).

Percentages of methylation (%CpG) were measured for each individual CpG present in the regions analysed by pyrosequencing. The regions chosen were around the CpGs identified by the Illumina methylation 450K analysis and include other CpGs. DNA methylation heatmaps were obtained using Prism8® software. The heatmaps in Figure 3 refer to the median of the median of the methylation percentages of the n CpG analysed by pyrosequencing (median (CpG1:%,…CpGn:%)). For each analysed gene, the difference of methylation percentages of gene promoter regions comparing WM266-4 and WM155, reported in Supplementary Figure 2A, was calculated as follows: median [(CpG1:%,…CpGn:%) in WM266-4 cells] - median [(CpG1:%,…CpGn:%) in WM115 cells].

### Definition of the signature score

The signature score considered the individual methylation values (percentages) of the selected single CpGs associated with *MYH1*, *PCDHB16*, *PCDHB15,* and the mean methylation values for the two CpGs selected for *BCL2L10*. For each gene, these methylation values were compared to the methylation median calculated from all the primary samples. A score of 1 was attributed to the gene when the methylation values differed by at least 15%. For *MYH1*, for which hypomethylation was associated with aggressiveness, this score was attributed when the methylation value was inferior to the median. Conversely, a score of 1 indicated a methylation value superior to the median for the three other genes (*PCDHB16*, *PCDHB15* and *BCL2L10*). The signature score was the sum of the scores attributed individually to the 4 genes and fell between 0 and 4. Signature score and survival information were reported in Supplementary Dataset3. This score was evaluated for each gene to assess potential correlation between individual gene scores. A random simulation using R software and based on a Chi-square test indicate that the methylation of these genes was independent from one another p< 0.01. Kaplan-Meier plots were created using GraphPad Prims8 software. The survival in months was indicated depending whether the score was under versus equal or superior than 2. Survival analysis was performed using a log-Rank and the Gehan-Breslow-Wilcoxon test with a p-value < 0.001 for each considered significant. Hazard ratio was estimated on R software using survival, survminer and ggplot2 packages.

### Methylation cluster identification through statistical analysis of the distribution of the hypermethylated genes

Identified methylation clusters highlight regions where the methylation distribution on the chromosome is not random. Maps of the 229 hypermethylated genes were visualized on the Ensembl website using the tool view on karyotype (http://www.ensembl.org). Sex chromosomes were excluded from the analysis. The clusters were defined as a group of at least two methylated genes in close proximity separated by non-methylated genes. The statistical relevance of the number of observed clusters on each chromosome was addressed using bootstrapping. For each simulation, methylated and non-methylated genes were randomly repositioned (shuffled) along each chromosome before recomputing the number of clusters. One thousand simulations were performed to estimate the probability of obtaining the number of observed clusters. All analyses were performed using custom-written scripts implemented in the statistical programming language R (http://cran.r-project.org/). All R- scripts are available from the authors upon request.

### Functional annotation and pathway analysis

The list of hypermethylated genes was imported into QIAGEN’s Ingenuity® Pathway Analysis (IPA®, QIAGEN Redwood City, www.qiagen.com/ingenuity). In IPA hypermethylated genes were mapped to molecular and cellular functions and to networks available in the ingenuity database and then ranked by score or p-value (p<0.05).

## RESULTS

### A three-step strategy identifies differentially methylated genes that identify melanoma aggressiveness

To identify genes whose DNA methylation state is related to the metastatic melanoma aggressiveness; we designed a strategy to compare the DNA methylome of three pairs of melanoma cell lines. Each pair was derived from the same patient melanoma cell lines that differed in aggressiveness and microenvironmental exposure with respect to their clinical origin or subsequent *in vivo* experimental processing (Figure 1). The first pair consisted of the WM115 and WM266-4 cell lines, derived from a vertical growth phase (VGP) primary melanoma and a cutaneous metastasis from the same patient, respectively, thus comparing a less and more aggressive pair of human melanoma cells. The second and the third pairs include a cell line established from a human lymph node metastasis (M4Be) and two metastatic variants selected for their increased metastatic potential in xenograft experiments either in mouse (M4BeS2) or in rat (TW12), (Figure 1A, step 1). It is important to note that in each pair, the aggressive cell line is derived from the same genetic background, but the most aggressive lines emerged in different *in vivo* contexts: human, mouse and rat, respectively.

**Figure 1.**
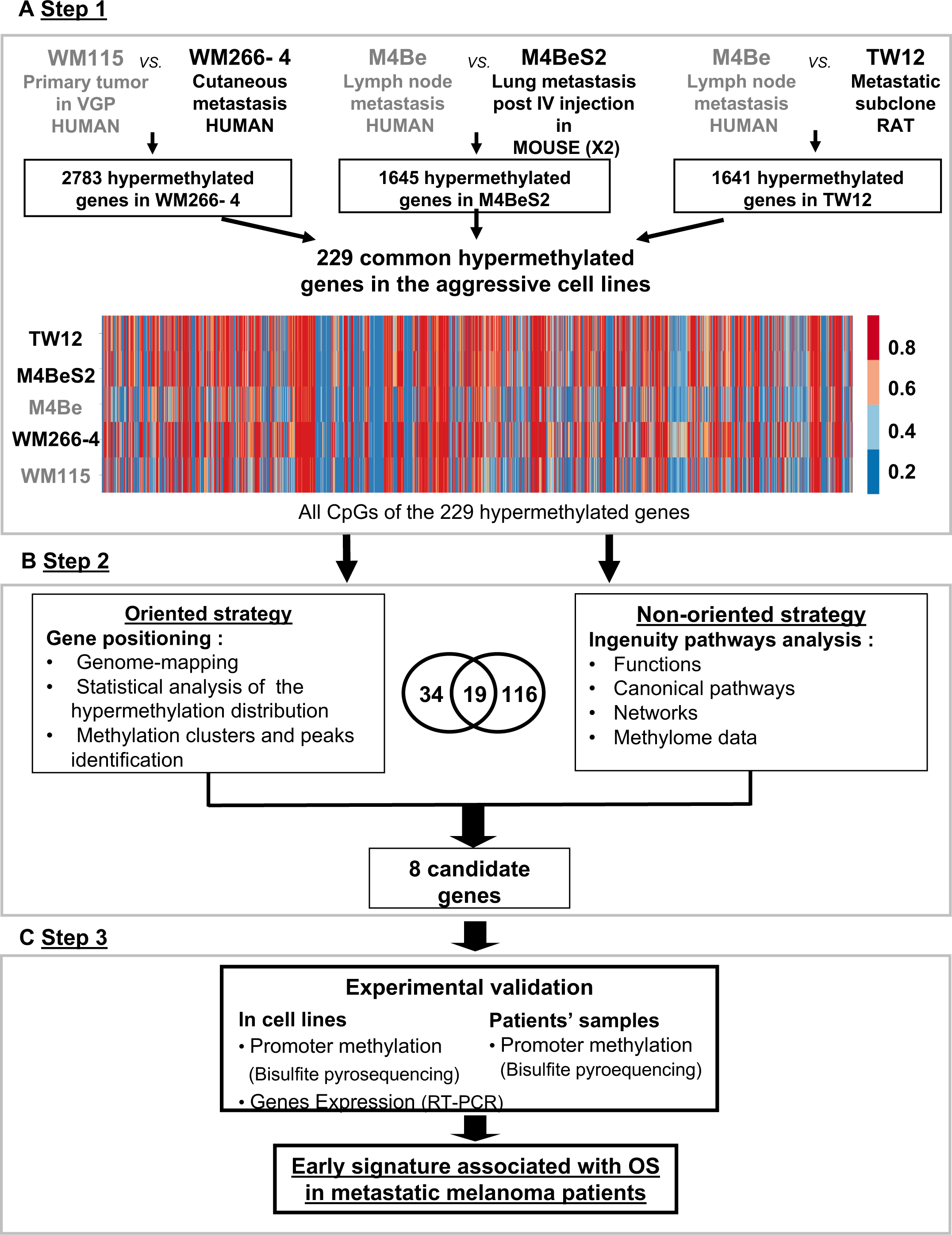
Strategy for identifying differentially methylated genes signatures of aggressive melanoma. The strategy is based on the analysis of three pairs of human melanoma cell lines with an aggressive variant derived under different physiological contexts: human, mouse and rat. In each pair, the cell line defined as more aggressive is indicated in bold characters. **A. Step1**: The methylation status of more than 480,000 CpG positions was compared in each cell line pair using the Illumina Infinium(®) Human Methylation 450K BeadChip technology. 229 common genes showing at least three CpGs positions with methylation levels increased by 20% in the aggressive cell line were retained (hypermethylated genes). **B. Step 2**: Two strategies for data analysis were used: the oriented strategy is based on a statistical analysis of the distribution of the hypermethylated genes across the genome, and the non-oriented strategy uses Ingenuity Pathway Analysis (IPA) software to identify potential links to described networks and functions. **C. Step3**: Experimental validation of the selected genes, by bisulfite pyrosequencing for DNA methylation and RT-PCR for gene expression, was performed in the WM115 and WM266-4 cell lines prior to analysis in patient samples. After applying this differential threshold to at least three CpG positions for each gene we found that 2783, 1645, and 1641 genes were hypermethylated in WM266-4 *vs.* WM115, M4BeS2 *vs.* M4Be, and TW12 *vs.* M4Be, respectively (A). 229 genes, comprising 5590 CpG sites, were common to all three pairs of cell lines. These 229 genes were further analysed using the human WM115/WM266-4 pair. 1287 (23%) CpGs were hypermethylated (>20%) in WM266-4 cells, of which 788 (61%) were located in promoter regions (TSS1500-TSS200-5’UTR-1^st^ exon), 452 (35%) in gene bodies and 47 (4%) in 3’UTR regions.

The DNA methylation profiles of each cell line was analysed using the Human Methylation 450K array BeadChip technology to identify the hypermethylated genes in the more aggressive variants. A first global analysis showed that nearly half of the analysed genes displayed at least one CpG position, where methylation levels are increased over 20% in the aggressive cell line compared to its respective counterpart. Following this first step of selection, we adopted two complementary approaches (Figure 1B). An oriented strategy, based on the genomic mapping of the 229 hypermethylated genes common to the aggressive melanoma cell lines, allowed us to identify clusters of hypermethylation. Clusters consisted of at least two hypermethylated genes that are either direct neighbours or separated within 3 megabases (Mb) of one another (Figure 2). Sex chromosomes were excluded from this analysis because they are subject to parental imprinting (20).

**Figure 2.**
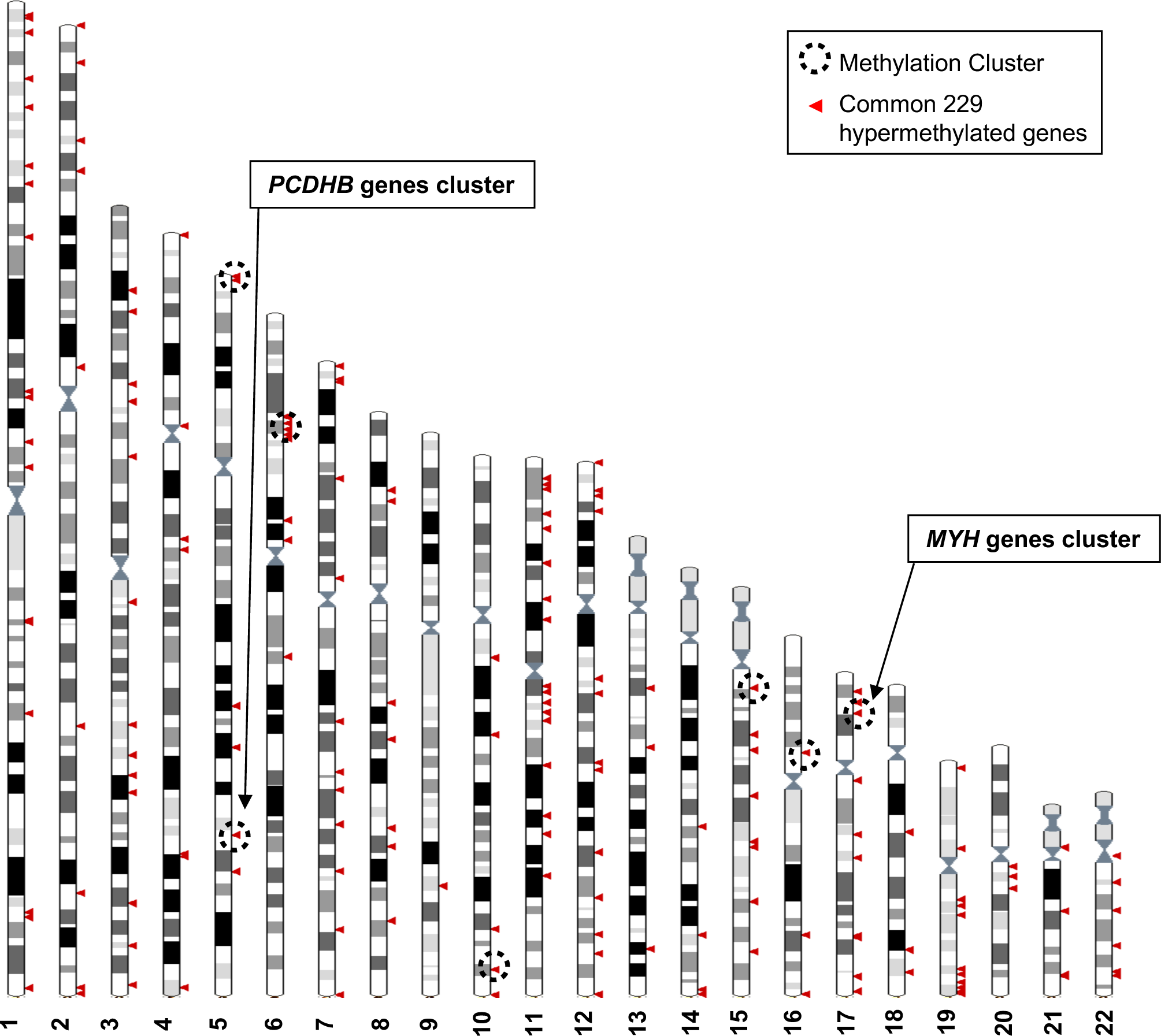
Genomic distribution of the 229 commonly hypermethylated genes in the more aggressive cell lines. The Ensembl genome browser (http://www.ensembl.org, view on karyotype) was used to map the 229 hypermethylated genes to the human genome. Sex chromosomes were excluded from the analysis. Each arrowhead could correspond to several genes. Methylation clusters are indicated by dotted line circles. Chromosome 5 and 6 circles correspond to two clusters that are too close to be separated on this scale.

Bootstrap analysis of the repartition of the 229 genes along the chromosomes confirmed a non-random distribution, hypermethylated genes enriched in short chromosomal regions (Figure 2 and Supplementary Figure S1). Nine methylation clusters were identified on chromosomes 5, 6, 10, 15, 16, and 17 (dotted line circles in Figure 2 and list in Supplementary Table S1), containing a total of 74 genes (Supplementary Table S2). Among these genes, 34 were further selected because they displayed at least two hypermethylated CpGs located in the promoter region (TSS1500-TSS200-5’UTR-1^ST^ exon) and a 40% difference in methylation when comparing human WM266-4 to WM115 cells. Chromosome 5 and 17 are of particular interest as the methylation clusters contain large multigenic families (Supplementary Figure S1). On chromosome 5, nine genes were identified as hypermethylated with a methylation cluster containing six genes belonging to the protocadherin beta (*PCDHB*) family (Supplementary Figure S1A and insert). *SPAG7*, *SOCS3* and *RAC3* displayed the strongest hypermethylation values (over 90%) among the 10 hypermethylated genes found on chromosome 17, that had at least two CpGs in the promoter region with a > 40% difference methylation in WM266-4 cells. Interestingly, this methylation cluster included five members of the multigenic Myosin heavy chain (MYH) family (insert in Supplementary Figure S1B).

The second strategy was non-oriented and consisted of analysing the functional pathways in which the 229 hypermethylated genes were involved. Using QIAGEN Ingenuity® Pathway Analysis (IPA®, QIAGEN Redwood City, www.qiagen.com/ingenuity) software, we found 116 genes highly associated with known functions (p < 0.05, Supplementary Table S3). The top functions, which might play a role in aggressiveness and carcinogenesis of the melanoma cells, were cell-to-cell signalling and interactions, cellular assembly and organization, cancer and cellular movement. Finally, when cross-checking the 34 genes from the oriented strategy and the 116 genes from the non-oriented strategy, we identified 19 common genes (Figure 1B). Notably, all of 19 genes were associated with one or two of the top functional networks from the IPA (Supplementary Dataset1).

### Gene selection and validation

The third part of our approach consisted of validating the methylation status of a subset of these candidate genes in melanoma cell lines and patient’s tissues samples. Combining the cluster analysis and the IPA results, we chose in total eight genes distributed on four different chromosomes, bearing hypermethylation clusters, and selected for representing an hypermethylation peak, for showing a strong methylation difference between the aggressive and non-aggressive cell lines, and for the potential role in aggressiveness suggested by the literature (Supplementary Table S4). On chromosome 17, we chose the following four genes. ***MYH1*** was selected because it forms a highly differentially methylated cluster, (Supplementary Figure S1B) and is known to show aberrant expression levels in aggressive cells in head and neck squamous and lung carcinoma tumours (21). ***SOCS3*** and ***RAC3*** displayed among the highest hypermethylation peaks in our aggressive melanoma lines (92% and 96%, respectively), with a very high differential methylation score, above 70%, between WM266-4 and WM115 cells. In addition, *SOCS3* has previously been reported to be hypermethylated in melanoma (22), while loss of *RAC3* expression has been associated with impaired invasion in glioma and breast carcinoma cells (23, 24). ***HOXB2*** was chosen for its lower methylation score (69%) and differential methylation score (45%). It has previously been associated with progression of bladder cancer when silenced by promoter hypermethylation, and it can be re-expressed upon demethylation treatment with 5-azacitidine (5azaC) (25). On chromosome 5, two genes were chosen from the PCDHB hypermethylation cluster: ***PCDHB15*** and ***PCDHB16*** that are CIMP-associated with bad prognosis in neuroblastoma (26, 27). Of note, neurons and melanocytes originate from the same germ layer during embryogenesis. Furthermore, ***BCL2L10*** (B Cell Lymphoma 2 like 10) located on chromosome 15 was selected because it bears the highest methylation score (73%) on this chromosome, it is implicated in apoptosis, it was previously described as hypermethylated in gastric cancer cell lines (28), and associated with poor prognosis in gastric cancer patients (29, 30). Finally, ***MIR155HG***, located on chromosome 21, encodes a microRNA, miR-155, was chosen because linked to cell proliferation and cancer (31), including in melanoma where it is downregulated (32, 33).

The methylation status of these eight genes (***MYH1, RAC3, SOCS3, HOXB2, PCDHB15, PCDHB16, BCL2L10, MIR155HG***) was further validated in WM115 *vs* WM266-4 cell lines by DNA pyrosequencing after bisulfite conversion and PCR amplification on 100 bp regions containing the CpGs identified in step 2 (Figure 1C). Seven genes pass the threshold of validation, 20% DNA methylation difference between the cell lines (Figure S2).

### Validation in patient samples and identification of a methylation signature

Next, we assessed the methylation profile of the eight selected genes in 20 tumour tissues from melanoma patients, of which 10 were from metastatic melanomas and 10 were from primary melanomas (Figure 1C). Four genes (*MYH1*, *PCDHB16*, *PCDHB15* and *BCL2L10*) showed a differential methylation profile between metastatic and primary tumour tissue samples (data not shown). For further validation, CpG sites in these four genes were individually analysed using bisulfite conversion followed by pyrosequencing in reference pair of cell lines (WM115/WM266-4) as well as two melanoma cell lines derived from the same patient: WM983A (primary site) and WM983B (lymph node metastatic site, Figure 3).

**Figure 3.**
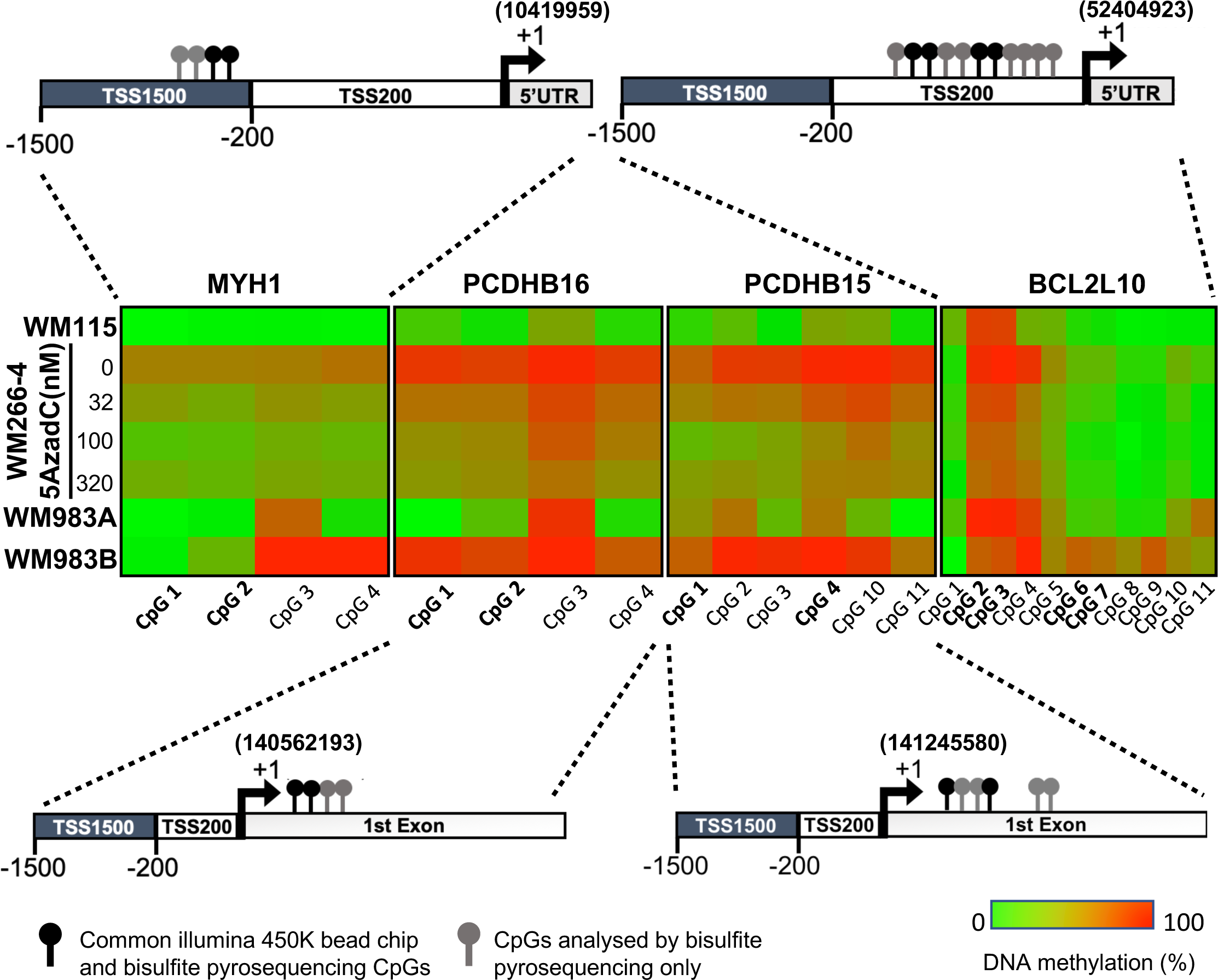
Promoter hypermethylation of the candidate genes in cell lines. **A.** Shown is the localization of all the CpGs analysed (lollipops) on the four selected genes (*MYH1*, *PCDHB15*, *PCDHB16*, and *BCL2L10*). The CpGs present on the 450k array are in bold type (black lollipops). The heatmap indicates the DNA methylation percentage (red = 100, green = 0) of the indicated CpGs in each gene and each cell line. WM115 and WM983A are primary cell lines derived from two patients. WM266-4 and WM983B are the cutaneous and lymph node metastasis counterpart. WM266-4 cells were treated with daily doses of 5AzadC (32, 100, and 320 nM) 72 hours before genomic DNA extraction.

DNA methylation levels at each CpG site were analysed by bisulfite pyrosequencing, confirming a clear difference in methylation for the four genes in the aggressive tumour cells compared to less aggressive forms. Interestingly, this DNA methylation could be reversed in WM266-4 cells using a low dose of 5azadC treatment (32, 100, and 320nM). The median methylation of these individual CpGs was determined on the first set of patient samples, 10 metastatic and 10 primary tumours, and on additional 10 primary tumour samples (Supplementary Figure S3). Remarkably, the median of DNA methylation within primary samples appeared to be inversely correlated with patient overall survival (OS). Thus, we defined patients with primary tumours diagnosis and long survival (LS) with an OS >1 year and patients with short survival (SS) with an overall survival (OS, median survival = 6 months) ≤1 year (median survival = 51 months) after diagnosed. The DNA methylation profile in primary tumours with SS similar to that observed in metastatic patients (top, in red, Supplementary Figure S3). The analysis showed that *MYH1* was globally hypomethylated in SS patients, whereas *PCDHB16*, *PCDHB15,* and *BCL2L10* were hypermethylated. Data from 29 additional primary tumours samples were then analysed. The extended cohort of patient with primary melanoma primary (n=49) confirmed that these four differentially methylated genes represent specific markers of more aggressive SS primary and metastatic melanoma tumours (Figure 4A).

**Figure 4.**
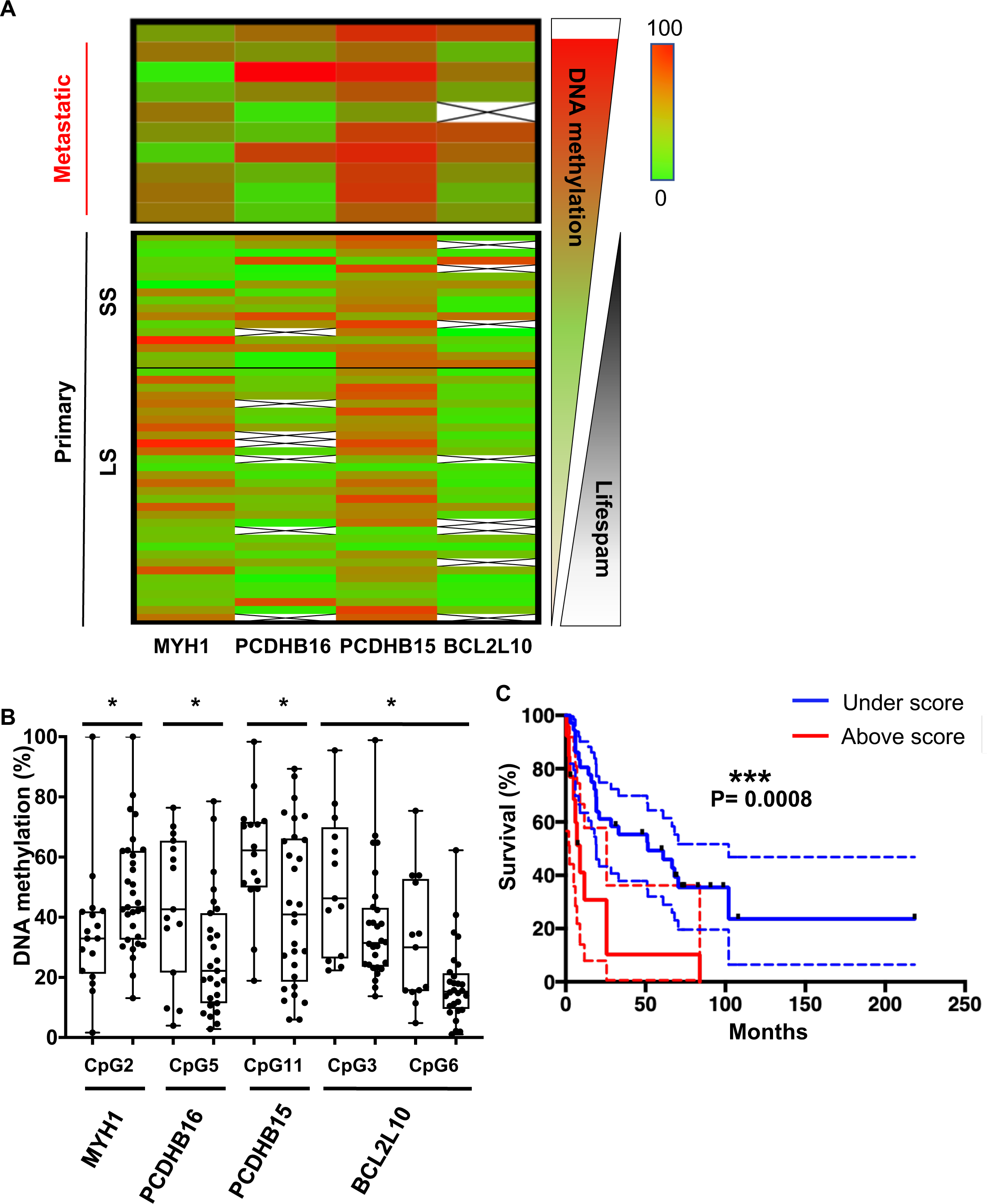
CpGs DNA methylation in primary melanomas predicts patient outcome. **A.** The heatmap shows the median DNA methylation of analysed CpGs for each gene in metastatic (n=10) and primary (n=49) patient samples. Primary samples are divided by a black line indicating the cut-off at one year between the short and long survival (SS (left) and LS right)). **B.** DNA methylation changes in selected CpGs between short and long survival patients (n=49). Fisher test to analyse variances and t-Test were performed, * = p< 0.05. **C.** Kaplan–Meier curve of patients’ survival based on the methylation score calculated for the five CpGs reported in panel B. The methylation score = 2 corresponds to at least two CpGs that showed a methylation difference >15% when compared to the DNA methylation median of the primary metastasis.

To determine the combination of CpGs that might predict melanoma aggressiveness, and thus survival outcome in patients with primary tumours, DNA methylation at individual CpG sites was analysed using bisulfite pyrosequencing (CpG positions are illustrated in Figure 3 and detailed in Supplementary Dataset2). One CpG showed significantly differential methylated in primary samples from SS patients (median SS = 6 months) when compared to LS patients (median LS = 51 months) for *PCDHB15*, *PCDBH16*, and *MYH1*, and two CpGs for *BCL2L10* (Figure 4B). DNA methylation *MYH1* CpG was inversely correlated with survival duration and was significantly hypomethylated in primary samples from SS patients; whereas *PCDHB16*, *PCDHB15,* and *BCL2L10* CpGs were hypermethylated in SS patients. To validate the robustness of this signature, a score was calculated as follows: a score of 1 was given to each gene when the median DNA methylation of the CpG met conditions for hypomethylation (>15%, seen for *MYH1*) and hypermethylation (>15%, for *PCDHB16*, *PCDHB15,* and *BCL2L10*). CpGs not meeting these criteria were given a score of 0. The final score was obtained by summing points attributed to each gene, so that scores range from 0 to 4 (Supplementary Dataset3). A threshold of 2 (*i.e.,* with at least two genes matching this condition) was used to include patients with a methylation score ≥ 2. The methylation score was represented as a function of survival in months (Figure 4C and Supplementary Figure S4A). Patients with a methylation score ≥ 2 (red line) had a shorter life expectancy (≤1 year) than those with a methylation score value <2 (blue line). This analysis demonstrated that a methylation score of at least 2 in primary melanoma samples is predictive of patient outcome (Log-rank test, p=0.0008) with a significant Hazard Ratio of 3.4 (p = 0.001, concordance index = 0.62, Supplementary Figure S4B). Then, we compared the methylation score to the clinical parameter used in clinic, the Breslow index. Based on the melanoma AJCC staging (T1, T2, T3 and T4), we confirmed that on our 49 patient samples an increased Breslow depth of the primary tumour is a prognostic factor for survival probability (Kaplan-Meier plot in Supplementary Figure S4C). As our cohort contained only one sample of T1 grade (Primary tumour’s depth less than 1mm), we then compared the survival probability between two groups: Breslow index below 2mm (T1 and T2) or above 2mm (T3 and T4). We obtained no statistical difference between the two groups (Supplementary Figures S4D and S4E), in contrast to what obtain upon use of the methylation signature (Supplementary Figures S4A and B). Most interestingly, the interaction between the methylation score and the Breslow index gave a significant increase of the hazard ratio from 3.4 to 6.3 (p < 0,001, concordance index = 0.63, Supplementary Figure S4F). This result support that the methylation score could improve the clinical prognostic utility of the Breslow index.

## DISCUSSION

To the best of our knowledge, the multistep strategy that we developed and used to identify differentially methylated genes in melanoma cells predicting aggressiveness is original. It is important to underline the starting point. We reasoned that, since epigenetic changes are linked to cell plasticity and biological environment changes (34–36), any common DNA methylation changes acquired by the most aggressive cells in different *in vivo* contexts (human, mouse and rat) would highlight a robust and relevant trait of melanoma tumour cell plasticity and aggressiveness despite their heterogeneity. Indeed, melanoma are highly heterogeneous tumours and until now it has been difficult to find a signature in primary melanoma, either based on DNA methylation or gene expression that help to predict prognosis as well as Breslow index (37). Several DNA methylation studies have been conducted comparing patient samples at different stages of cutaneous melanoma to normal samples *,i.e.,* melanocytes or nevi (38–41) or using cell lines derived from multi-grade patient samples (42) (9). While such studies have identified genes regulated by DNA methylation, none have yet identified a common pattern or a specific signature of melanoma aggressiveness. In contrast, the unique approach used in our study yielded a potential DNA methylation signature that correlates with outcomes.

Considering that the cellular models utilized, have experienced very different experimental microenvironments, we assumed that the common 229 identified hypermethylated genes represented genes either playing a direct role in or being associated with the aggressiveness of melanoma. Next, we applied a bootstrap analysis of the distribution of these hypermethylated genes revealing that they are not randomly distributed along the chromosomes, but are instead organized in clusters of hypermethylated genes. This observation of clustered genes regulated by DNA methylation is reminiscent of the mechanism underlying parental imprinting, and its spreading on imprinted genes during development (43, 44). It is also coherent with the association of hypermethylated regions with long range epigenetic silencing (LRES) described by Frigola J. *et al* (45) in colon cancer, and with the fact that LRES regions containing hypermethylated genes extend up to 4 Mb and are correlated with gene extinction (46). Thus, guided by the hypothesis that these clustered genes correspond to early changes in CpG methylation during tumour progression, and may thus constitute a starting point for spreading of DNA methylation, we concentrated our efforts on these. An IPA analysis of the corresponding biological functions related to tumorigenesis or resistance to therapy, facilitated selection of eight potential genes that could be early signals of the aggressiveness of the disease. The DNA methylation signature at these eight genes was analysed in additional melanoma cell lines (WM983A and WM983B) derived from the same patient, and in patient metastatic and primary tumour samples, highlighting four genes of interest. All four genes are demethylated upon treatment with 5AzadC in WM266-4 metastatic melanoma cells, and displayed the strongest methylation differential.

In the last part of our study, we challenged the methylation status of these four gene promoters in 59 patient samples, Surprisingly, five CpGs were significantly differentially methylated between patients with a short overall survival (<1-year, SS) and those with a longer overall survival (>1 year, LS). This led to the discovery of an early and robust signature of melanoma progression that is based directly on the primary tumour that can significantly predict patient outcome (p=0.0008). Several studies have described different methylation patterns at various stages of melanoma disease, but only one reported a difference in methylation profile among primary tumour samples linking it to an ulceration status and thus a poor clinical outcome (47). More recently, Guo *et al.* (8) identified four DNA methylation biomarkers by analysing all melanoma types in the TCGA database, but without validating this on another patient cohort. Importantly, we have discovered CpG sites and genes that are key to metastatic melanoma formation and are grouped in genomic clusters. Our signature is unique, as is the integrated approach and the baseline assumption used to identify it. Going forward, a signature could be used to develop a much-needed early diagnostic DNA methylation kit, similar to existing kits for colon, lung and prostate cancer (48–50).

## CONCLUSIONS

We developed a novel multistep approach that allowed us to identify a methylation signature of five CpGs in primary melanoma tissues that has the potential to predict survival outcomes in cutaneous melanoma patients. Our method was based on two main concepts, the first one being that aggressive traits marked by DNA hypermethylation appear early in the disease and are independent of physiological context. The second concept is that hypermethylated sites in metastatic forms of melanoma are gathered in genomic clusters. On the methodological side, we combined analysis of the DNA methylome with chromosomal location. Following these general concepts, this integrated approach can be applied not only to other cancer types, but also to other diseases or biological processes such as aging and development.

## ABBREVIATIONS

5AzadC: 5aza-2′-deoxycytidine
AJCC: American joint committee on cancer CIMP: CpG island methylator phenotype
CpG: Cytosine preceeding guanine nucleotide dimer 5’-3’direction
DNA: Deoxyribonucleic acid
FFPE: formalin-fixed paraffin-embedded
IPA: Ingenuity pathway analysis
LRES: Long range epigenetic silencing LS: long survival
Mb: Megabases
OVS: overall survival
PCR: Polymerase chain reaction
RNA: ribonucleic acid
RRBS: Reduced representation bisulfite sequencing
SS: short survival
TCGA: The cancer genome atlas
TSS: Transcription start site VGP: Vertical growth phase

## DATA AVAILABILITY

R-scripts are available upon request to the authors.

The DNA methylome dataset has been deposited in NCBI’s Gene Expression Omnibus under the GEO Series accession number GSE155856 (https://www.ncbi.nlm.nih.gov/geo/query/acc.cgi?acc=GSE155856)

The datasets supporting the conclusions of this article are included within the article and the following Supplementary files.

## SUPPLEMENTAL MATERIALS

Supplementary file1.pdf: Supplementary figures and tables Supplementary_Dataset1.xls: IPA software networks results.

Supplementary_Dataset2.xls: Bisulphite pyrosequencing primer, sequences, and CpG location.

Supplementary_Dataset3.xlsx: Survival data and signature score for each patient sample.

## ACKNOWELDGMENTS

Not applicable.

## Supplementary Information

**Fig. S1.**
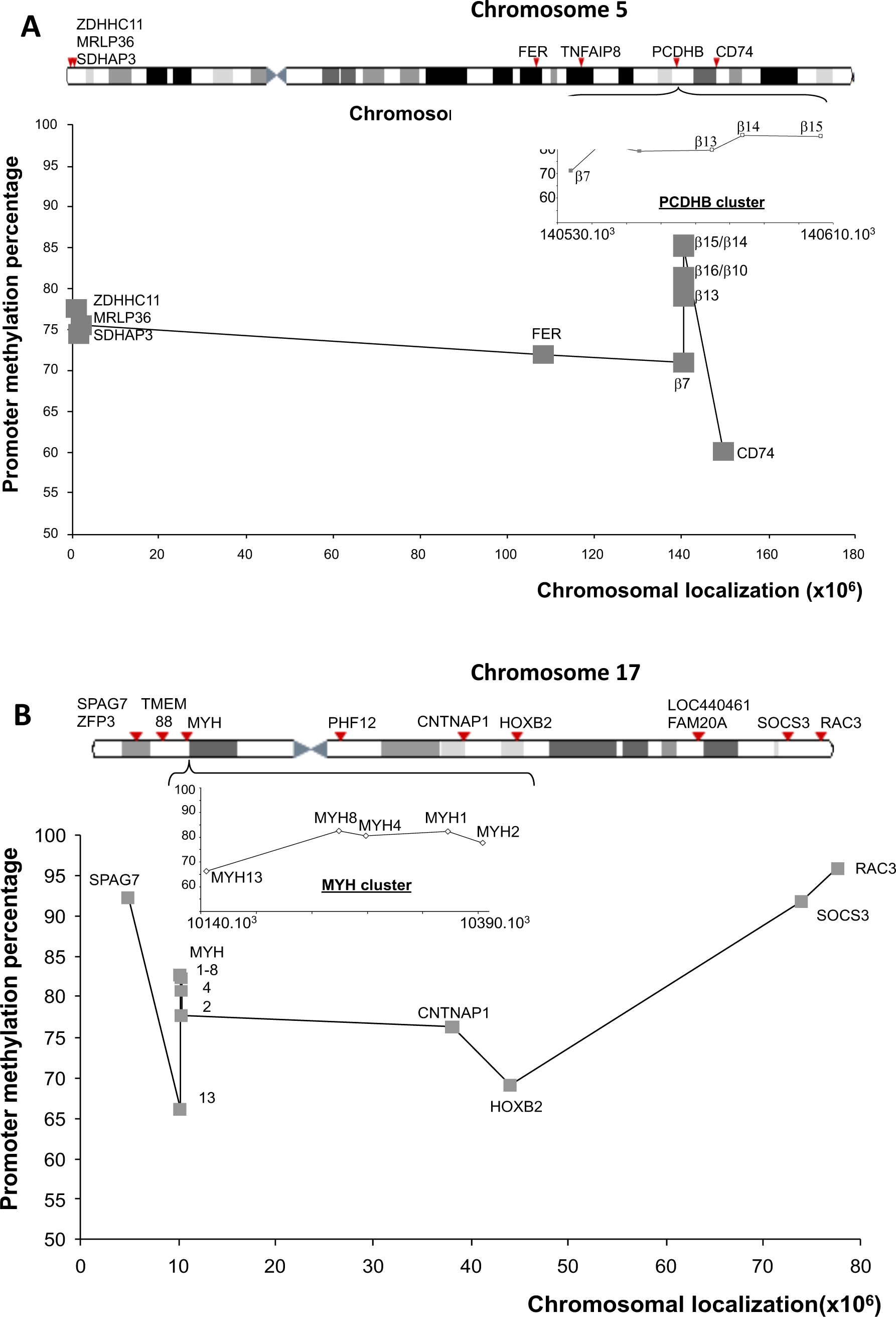
Examples of genomic localization of the genes hypermethylated in the most aggressive cell lines. Promoter methylation scores correspond to the average methylation values at CpG positions located in the promoter regions showing an increased methylation above 20% in WM266-4 compared to WM115 cells. The genes are indicated on the “band giemsa-related” representation (arrows). The graph shows genes with at least two CpGs in the promoter region with DNA methylation differences over 40% between WM266-4 and WM115 cells (grey squares). A) Localization and promoter methylation score of nine hypermethylated genes found on chromosome 5. The PCDHβ genes cluster is magnified in the insert. B) Localization and promoter methylation score of 15 hypermethylated genes found on chromosome 17; five of them belong to the MYH1 cluster (insert).

**Fig. S2.**
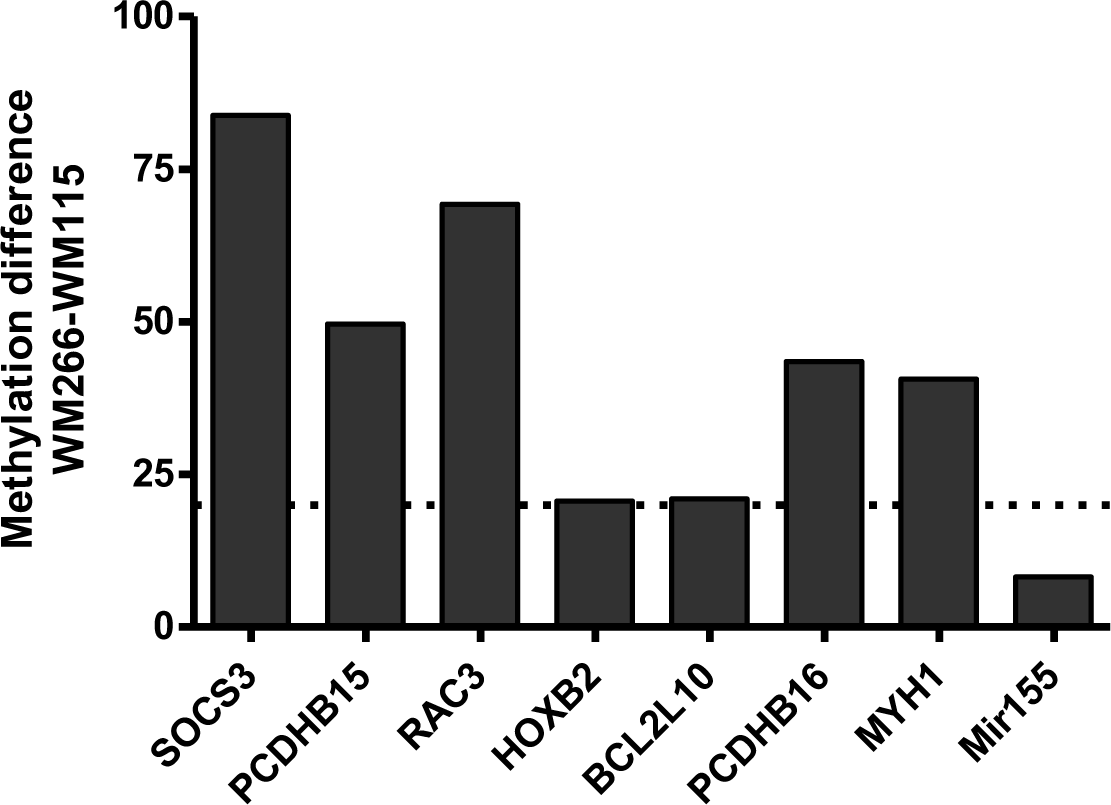
Promoter hypermethylation of the candidate genes in WM266-4 *vs* WM115 cells. Hypermethylation (Methylation difference ≥ 20%: dotted line) for all candidate genes analysed by Bisulfde-Pyrosequencing.

**Fig. S3.**
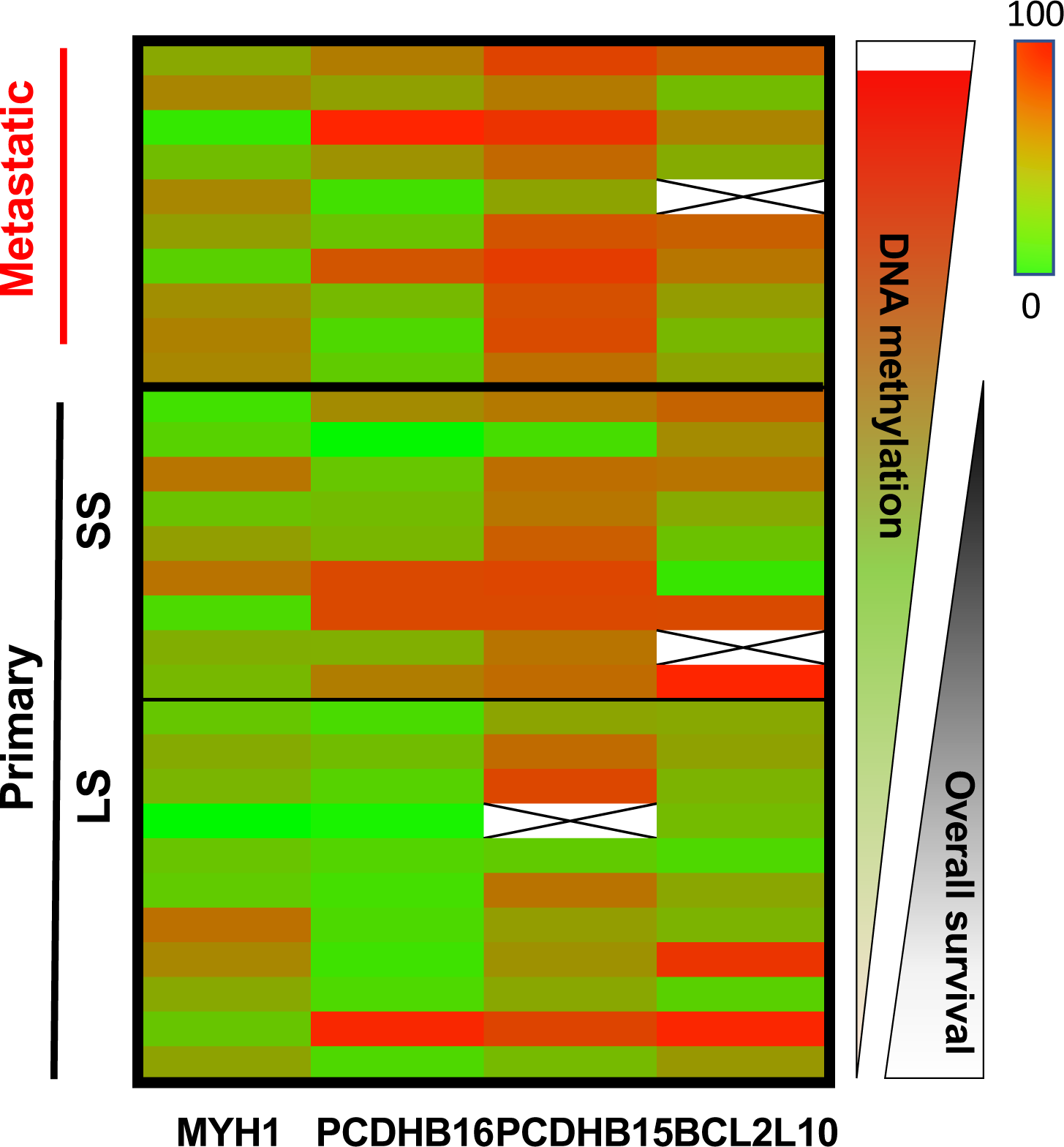
Heatmap of DNA methylation of MYH1, PCDHB16, PCDHB15 and BCL2L10 in the first set of 10 metastatic melanoma patient sample and 20 primary patient melanoma. The median DNA methylation of the analysed CpG for each gene is indicated as percentage and overall survival in months. Primary samples are divided by a small black line indicating the cut-off at one year between the short and long overall survival (SS and LS).

**Fig. S4.**
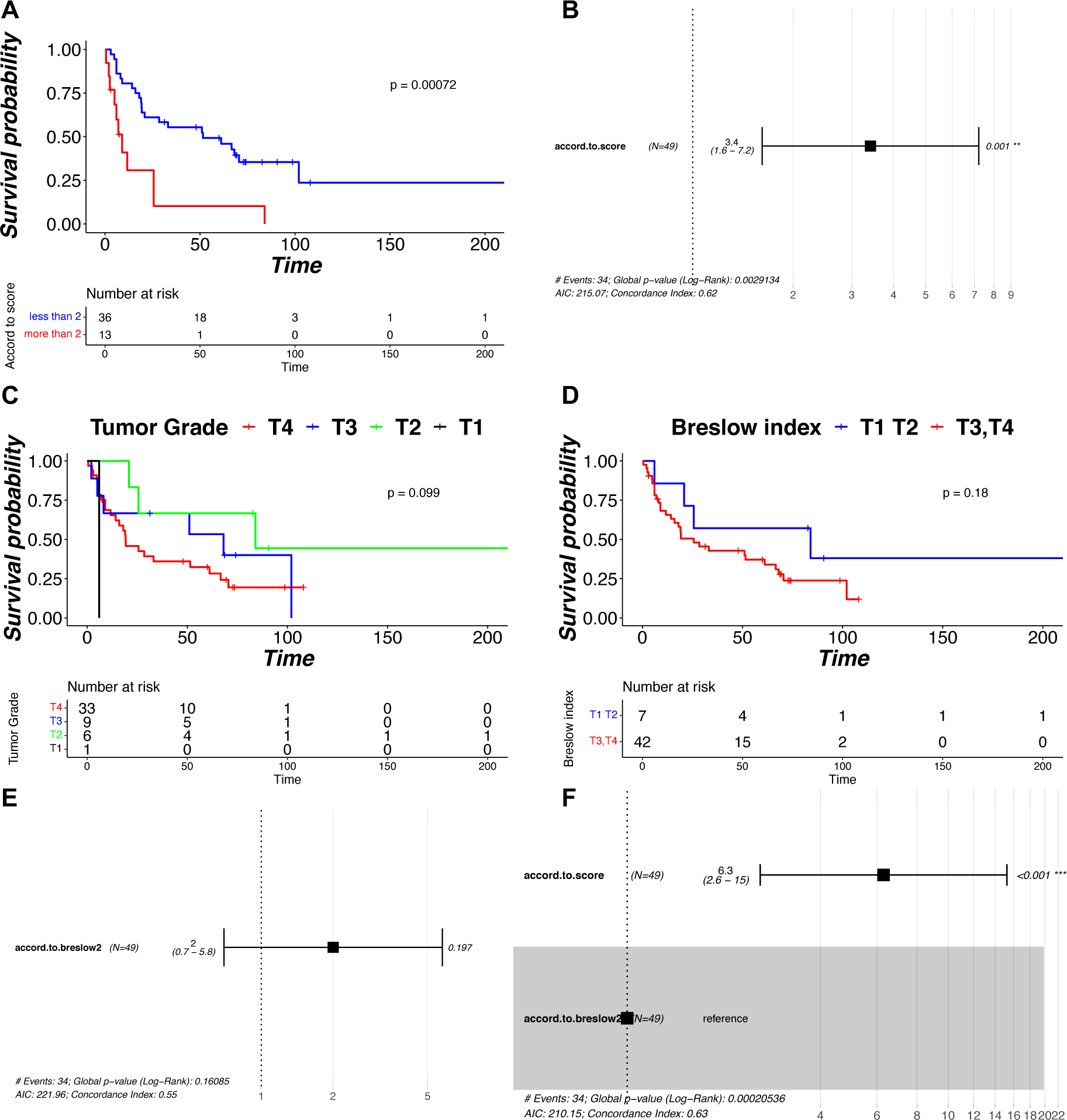
The DNA methylation score is a better prognostic factor than classical used clinical parameter Breslow index. A) Kaplan–Meier curve of patients based on the methylation score in the 49 primary melanoma samples. B) Hazard ratio plot calculated from our methylation score on the 49 primary melanoma samples. C) Kaplan–Meier curve of patients according to the primary tumor grade defined by the AJCC staging system of melanoma from Breslow index (Primary tumor T1 = less than 1mm, T2 = less than 2 mm, T3 = between 2 and 4 mm, T4 = more than 4 mm). D) Kaplan–Meier curve of patients based on the Breslow index in the 49 primary melanoma samples (less than 2 in blue: T1 and T2 grade, more than 2 in red: T3 and T4 grades). E) Hazard ratio plot calculated from Breslow index on the 49 primary melanoma samples. F) Hazard ratio plot calculated from the interaction between our methylation score and the Breslow index on the 49 primary melanoma samples.

**Table S1.** Clusters of hypermethylated genes identified by the oriented strategy. Nine methylation clusters containing 29 genes on six chromosomes.

| Cluster | Gene Name | Ensembl Gene ID | EntrezGene ID | chromosome | Ref Seq Genes Position |  |
| --- | --- | --- | --- | --- | --- | --- |
|  |  |  |  |  | Start (bp) | end (bp) |
| 1 | SDHAP3 | ENSG00000185986 | 728609 | 5 | 1572073 | 1594646 |
|  | MRPL36 | ENSG00000171421 | 64979 | 5 | 1798499 | 1799956 |
| 2 | PCDHB16 | ENSG00000196963 | 57717 | 5 | 140561265 | 140565796 |
|  | PCDHB9 | ENSG00000120324 | 56127 | 5 | 140566893 | 140571111 |
| 3 | PCDHB13 | ENSG00000187372 | 56123 | 5 | 140593491 | 140596993 |
|  | PCDHB14 | ENSG00000120327 | 56122 | 5 | 140601898 | 140605860 |
|  | PCDHB18 | ENSG00000146001 | 54660 | 5 | 140613938 | 140617101 |
|  | PCDHB15 | ENSG00000113248 | 56121 | 5 | 140624917 | 140627801 |
| 4 | KIAA1949/PPP1R18 | ENSG00000146112 | 170954 | 6 | 30644166 | 30655672 |
|  | NRM | ENSG00000137404 | 11270 | 6 | 30655824 | 30659197 |
|  | MDC1 | ENSG00000137337 | 9656 | 6 | 30667584 | 30685458 |
| 5 | BAT4/GPANK1 | ENSG00000204438 | 7918 | 6 | 31629006 | 31633407 |
|  | CSNK2B | ENSG00000204435 | 1460 | 6 | 31633657 | 31637843 |
| 6 | DOCK1 | ENSG00000150760 | 1793 | 10 | 128594023 | 129250780 |
|  | FAM196A | ENSG00000188916 | 642938 | 10 | 128933690 | 128994422 |
| 7 | SNORD115-15 | ENSG00000201679 | 100033453 | 15 | 25451409 | 25477615 |
|  | SNORD115-21 | ENSG00000199833 | 100033603 | 15 | 25453230 | 25453310 |
|  | SNORD115-42 | ENSG00000201143 | 100033816 | 15 | 25492492 | 25492573 |
| 8 | BOLA2 | ENSG00000183336 | 552900 | 16 | 29464914 | 29466285 |
|  | BOLA2B | ENSG00000169627 | 654483 | 16 | 29464914 | 29466285 |
|  | GIYD1/SLX1A | ENSG00000132207 | 548593 | 16 | 29465822 | 29469545 |
|  | GIYD2 | ENSG00000181625 | 79008 | 16 | 29465822 | 29469545 |
|  | SULT1A3 | ENSG00000261052 | 6818 | 16 | 29471207 | 29476301 |
|  | SULT1A4 | ENSG00000213648 | 445329 | 16 | 29471207 | 29476301 |
| 9 | MYH13 | ENSG00000006788 | 8735 | 17 | 10204183 | 10276322 |
|  | MYH8 | ENSG00000133020 | 4626 | 17 | 10293642 | 10325267 |
|  | MYH4 | ENSG00000264424 | 4622 | 17 | 10346608 | 10372876 |
|  | MYH1 | ENSG00000109061 | 4619 | 17 | 10395627 | 10421859 |
|  | MYH2 | ENSG00000125414 | 4620 | 17 | 10424465 | 10453017 |

**Table S2:**
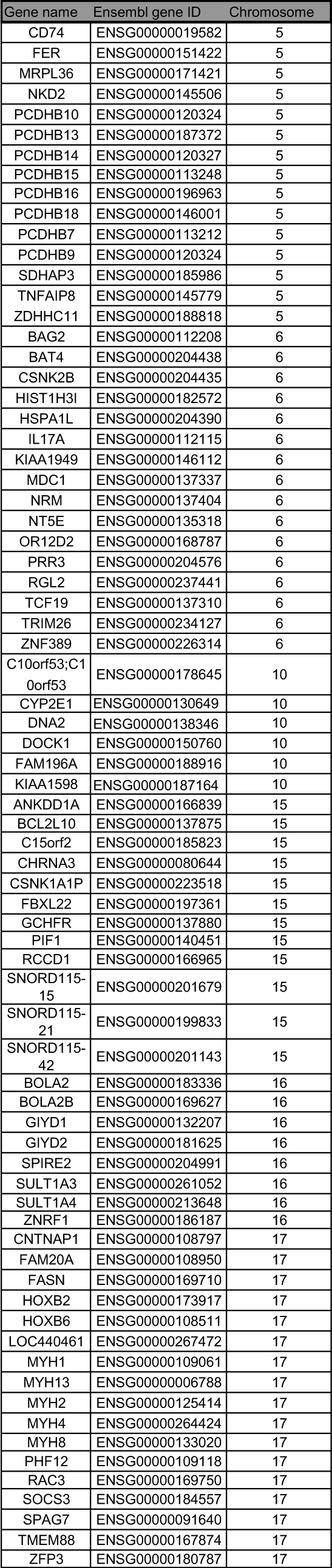
List of the 74 hypermethylated genes found on six chromosomes bearing at least one cluster.

**Table S3:**
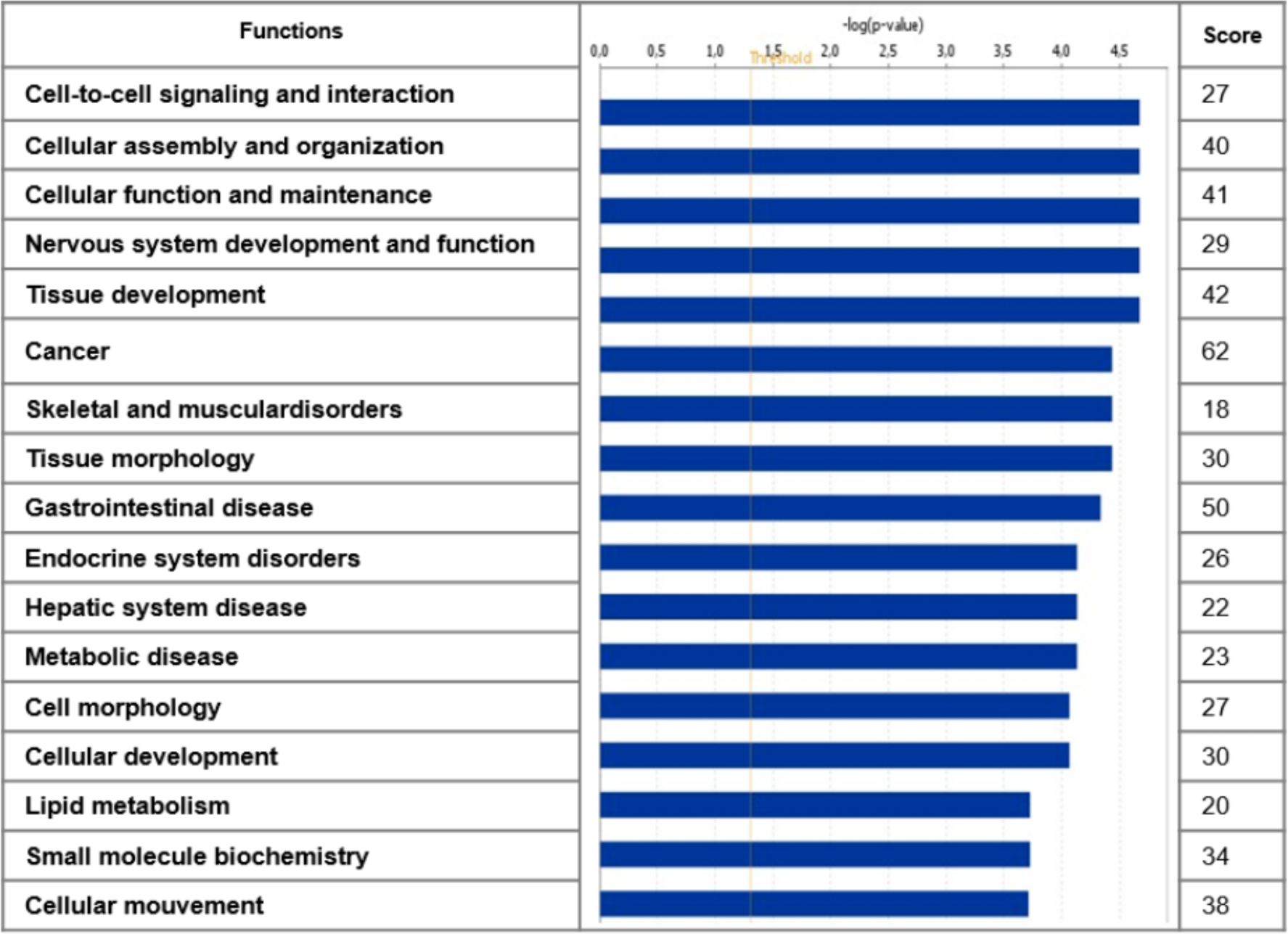
Top Functions associated by IPA to the 229 hypermethylated genes in the aggressive cell lines. Among the 229 genes, 116 were associated to IPA canonical functions. Column 1 indicates the top functions. Column 2 shows the -log(p-value) as horizontal histogram relative to the Fisher exact test (p < 0.05 : -log (0.05 = 1.3 =threshold) used by IPA to classify the most representative functions. The score in column 3 corresponds to the number of genes among the 229 belonging to the associated function.

| Functions | $-\log(p\text{-value})$ | Score |
| --- | --- | --- |
| Cell-to-cell signaling and interaction |  | 27 |
| Cellular assembly and organization |  | 40 |
| Cellular function and maintenance |  | 41 |
| Nervous system development and function |  | 29 |
| Tissue development |  | 42 |
| Cancer |  | 62 |
| Skeletal and muscular disorders |  | 18 |
| Tissue morphology |  | 30 |
| Gastrointestinal disease |  | 50 |
| Endocrine system disorders |  | 26 |
| Hepatic system disease |  | 22 |
| Metabolic disease |  | 23 |
| Cell morphology |  | 27 |
| Cellular development |  | 30 |
| Lipid metabolism |  | 20 |
| Small molecule biochemistry |  | 34 |
| Cellular movement |  | 38 |

**Table S4:** Detailed informations of the eight selected genes.

| Gene symbol | Gene | RefSeqGene | Chromosome | Methylation location | $\Delta$ methylation (%)<br>(βWM266-4 - βWM115)<br>x 100 | WM266-4 cells<br>promotor methylation (%) | IPA Function |
| --- | --- | --- | --- | --- | --- | --- | --- |
| MYH1 | myosin, heavy chain 1 | NM_005963 | 17 | cluster | 75 | 82 | Cellular assembly and organization, cancer |
| SOCS3 | suppressor of cytokine signaling 3 | NM_003955 |  | Methylation peak | 72 | 92 | Cancer |
| RAC3 | RAS-related C3 botulinum toxin substrate 3 | NM_005052 |  | Methylation peak | 70 | 96 | Cellular assembly and organization, cancer, cellular movement |
| HOXB2 | homeobox B2 | NM_002145 |  | chromosome | 45 | 69 | Tissue development |
| PCDHB15 | protocadherin beta 15 | NM_018935 | 5 | cluster | 45 | 85 | cell-to-cell signaling and interaction |
| PCDHB16 | protocadherin beta 16 | NM_020957 |  | cluster | 52 | 82 | cell-to-cell signaling and interaction |
| BCL2L10 | BCL2-like 10 | NM_020396 | 15 | chromosome | 46 | 73 | Cancer |
| Mir-155HG | Mir-155 Host Gene | NR_001458 | 21 | other | 32 | 74 | Cancer |

**Table S5.**
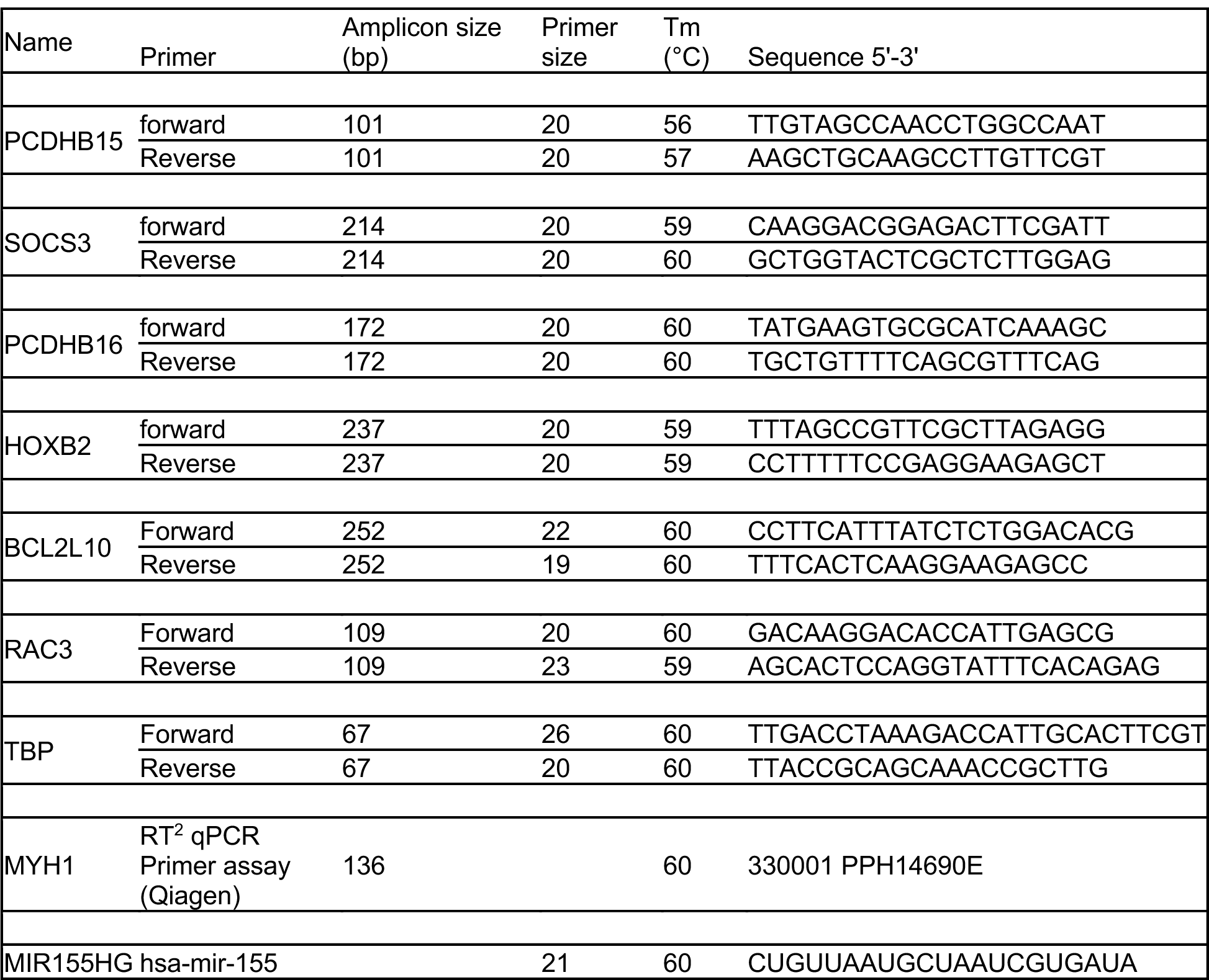
Primers informations for RT-PCR and RT-qPCR experiments.

**Dataset S1 -** Results from the IPA software networks.

**Dataset S2 –** Gene sequences, Illuminina CpGs and bisulfite pyrosequencing primers

**Dataset S3 –** Survival data and signature score for each patient sample.

